# Impaired H19 lncRNA expression contributes to the compromised developmental angiogenesis in EVL-deficient mice

**DOI:** 10.1101/2023.04.19.537575

**Authors:** Joana Zink, Timo Frömel, Reinier A. Boon, Ingrid Fleming, Peter M. Benz

## Abstract

Endothelial tip cells are essential for VEGF-induced angiogenesis, but underlying mechanisms are elusive. Endothelial-specific deletion of EVL, a member of the mammalian Ena/VASP protein family, reduced the expression of the tip cell marker protein endothelial cell specific molecule-1 (Esm1) and compromised the radial sprouting of the vascular plexus in the postnatal mouse retina. The latter effects could at least partly be attributed to reduced VEGF receptor 2 (VEGFR2) internalization and signaling but the underlying mechanisms(s) are not fully understood. In the present study, we revealed that the expression of the long non-coding RNA H19 was significantly reduced in endothelial cells from postnatal EVL^-/-^ mice and in siRNA-transfected human endothelial cells under hypoxic conditions. H19 was recently shown to promote VEGF expression and bioavailability via Esm1 and hypoxia inducible factor 1α (HIF-1α). Similar to EVL^-/-^ mice, the radial outgrowth of the vascular plexus was significantly delayed in the postnatal retina of H19^-/-^ mice. In summary, our data suggests that loss of EVL not only impairs VEGFR2 internalization and downstream signaling, but also impairs VEGF expression and bioavailability in the hypoxic retina via downregulation of lncRNA H19.

## Introduction

In the angiogenic vasculature, highly motile and invasive endothelial tip cells form actin-rich lamellipodia and filopodia, which probe the environment for guidance cues, such as vascular endothelial growth factor (VEGF), and thereby determine the direction of growth (Carmeliet et al., 2009). VEGF binding triggers the phosphorylation and internalization of endothelial VEGF receptor 2 (VEGFR2), which is crucial for the activation of downstream signaling targets that control angiogenic sprouting (Sakurai et al., 2005; Simons et al., 2016). In mammals, the Ena/VASP family of proteins consists of mammalian enabled (Mena), VASP, and Ena-VASP-like protein (EVL). These Ena/VASP proteins are important mediators of cytoskeletal control, linking kinase signaling pathways to actin assembly (Faix and Rottner, 2022). The proteins regulate many cardiovascular processes, including platelet adhesion and aggregation, endothelial barrier function, heart contraction, and vascular repair after ischemia (Aszodi et al., 1999; Benz et al., 2008; Benz et al., 2013; Laban et al., 2018; Massberg et al., 2004). More recently, Ena/VASP proteins have also been implicated in the regulation of developmental angiogenesis in vivo (Benz et al., 2019; Zink et al., 2021) and EVL deletion resulted in a significantly delayed radial sprouting of the vascular plexus in postnatal retina. The underlying mechanism involves impaired VEGFR2 internalization and signaling but is not fully understood (Zink et al., 2021).

The human genome is extensively transcribed and give rise to thousands of long non-coding RNAs (lncRNAs), which are not translated into functional proteins but play a critical role in epigenetic gene regulation (Busscher et al., 2022). The lncRNA H19 was one of the first to be discovered. H19 is enriched in the cardiovascular tissue and dysregulation of H19 has been linked to the development of many severe cardiovascular diseases, including diabetic retinopathy, atherosclerosis, cardiac diseases, pulmonary hypertension and others (Busscher et al., 2022). However, H19 has also been implicated in endothelial cell aging via inhibition of STAT3 signaling (Hofmann et al., 2019) as well as cancer progression by activating epithelial-mesenchymal transition, the cell cycle and cancer angiogenesis via mechanisms like microRNA (miRNA) sponging – the binding to and inhibition of miRNA activity (Shermane Lim et al., 2021). However, a role of H19 in developmental angiogenesis is currently unknown.

The aim of the present study was to assess a possible link between EVL and the lncRNA H19, which may contribute to the impaired developmental angiogenesis in EVL-deficient mice.

## Results and Discussion

To better understand the molecular mechanisms underlying the impaired developmental angiogenesis in EVL^-/-^ mice, the gene expression profiles of CD31 and CD34 double-positive endothelial cells from wild-type and EVL^-/-^ mouse retinas on postnatal day 5 (P5) were compared by RNA sequencing (Zink et al., 2021) and (Figure 1). Fitting with impaired VEGF-signaling, gene set enrichment analysis of EVL- deficient endothelial cells revealed significant changes in angiogenesis-related gene ontology terms, including blood vessel morphogenesis and endothelium development. Protein coding genes that were markedly downregulated in EVL^-/-^ endothelial cells included the tip cell marker Esm1, which is well known to promote angiogenic sprouting (Rocha et al., 2014). Consistent with our transcriptome data, retinal Esm1 protein expression and tip cell numbers were significantly reduced at the vascular front of P5 EVL^-/-^ retinas (Zink et al., 2021). One of most strongly and significantly changed non-coding transcripts was lncRNA H19. Compared to wild-type controls, H19 expression was reduced by approximately 65% in EVL-deficient retinal endothelial cells (Figure 2A). To confirm the impact of EVL- deficiency on H19 RNA levels, EVL was downregulated (siRNA) in human umbilical vein endothelial cells. In EVL^-/-^ mice, retinal angiogenesis was particularly impaired in the early postnatal days, when the retina is still largely hypoxic (Zink et al., 2021). To mimic this situation in vitro, we exposed the endothelial cells to normoxia (room air) or hypoxia (1% O_2_) and analyzed EVL and H19 expression by qPCR. Irrespective of the oxygen concentration, gene-specific siRNA treatment reduced EVL RNA levels to approximately 20% of control siRNA transfected cells (Figure 2B). Under normoxic conditions, H19 levels were slightly decreased in EVL siRNA treated cells, but this difference was not significant. Previously, hypoxia has been shown to induce H19 expression through direct and indirect Hif-1α activity (Wu et al., 2017). Consistent with this study, H19 levels were significantly increased in both control and EVL siRNA-transfected cells under hypoxic conditions. However, compared to control cells, H19 levels were significantly decreased in endothelial cells with siRNA-mediated knockdown of EVL (Figure 2C), indicating that EVL deficiency impairs H19 expression particularly under hypoxic conditions. Given that the P5 mouse retina is still partially hypoxic (West et al., 2005), this finding is also in good agreement with the reduced expression of H19 in the sorted retinal endothelial cells. Given its established role in cancer angiogenesis, we determined whether or not the knockout of H19 had an impact on postnatal retinal angiogenesis in mice. Relative to littermate controls, the radial outgrowth of the vascular plexus from the optic nerve head to the retinal periphery at P5 was significantly delayed in H19-deficient mice. The magnitude of the delay was similar to the vascular defect observed in global EVL knockout mice (Figure 3A, B). Therefore, reduced H19 expression in EVL- deficient mice may constitute an additional link to the observed vascular phenotype. Fitting with this hypothesis and the reduced Esm1 levels in EVL-deficient animals, impaired lncRNA H19 expression has recently been shown to downregulate Esm1 expression via reduced sponging of microRNA 181b-5p (Li et al., 2022). Esm1 in turn has been shown to enhance VEGF bioavailability and thereby increase retinal angiogenesis (Rocha et al., 2014), indicating that H19 deficiency reduces VEGF levels via Esm1. However, there is also an Esm1-independent link to VEGF signaling. Hypoxia is the primary driver of VEGF release and VEGF-induced neovascularization in the postnatal mouse retina (Liu et al., 1995; Stone et al., 1995) and H19 was recently shown to promote VEGF expression via hypoxia inducible factor 1α (HIF-1α) in glioma cells (Liu et al., 2020). In summary, our data suggests that loss of EVL not only impairs VEGFR2 internalization and downstream signaling, but also impairs VEGF expression and bioavailability in the hypoxic retina via downregulation of lncRNA H19.

**Figure 1.**
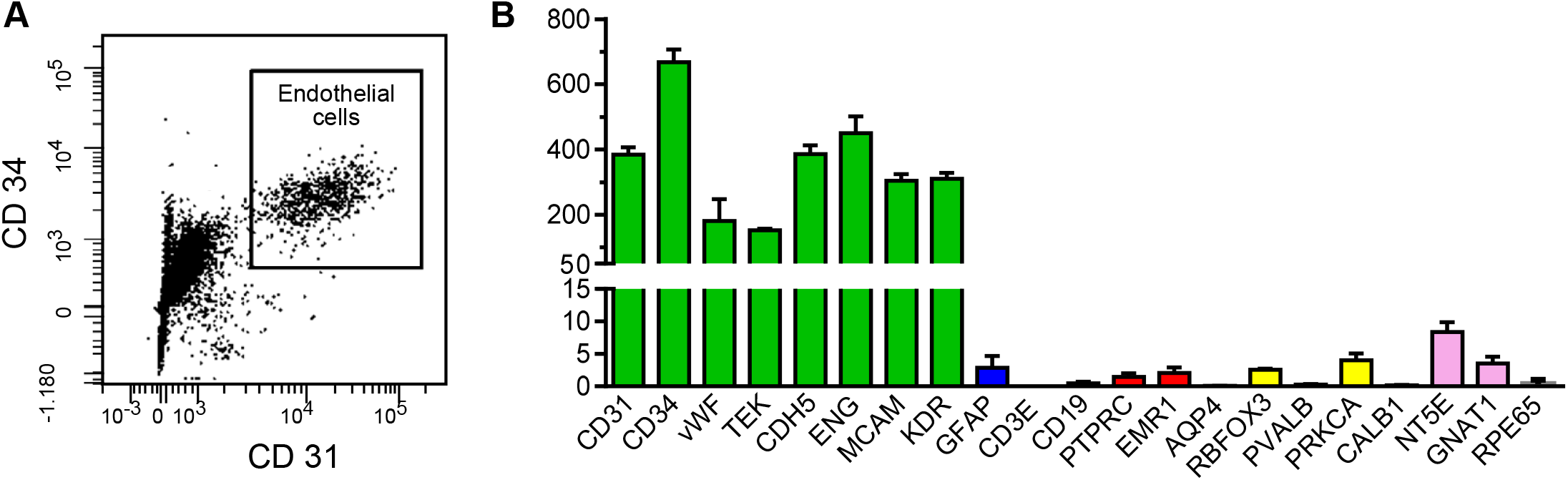
RNA sequencing of wild-type and EVL^-/-^ P5 retinal endothelial cells. (**A**) Postnatal day 5 (P5) retinas from wild-type and EVL^-/-^ mice were digested with collagenase and dispase. CD31 and CD34 double-positive endothelial cells were isolated by FACS and analyzed by RNA-sequencing on an Illumina HiSeq 4000 sequencer generating 50 bp single-end reads (ca. 30-40 Mio reads/sample). (**B**) RNA sequencing of CD31 and CD34 double-positive retinal endothelial cells from P5 wild-type mice. RNA levels (FPKM; fragments per kilobase million) of marker genes of endothelial cells (green; CD31 (gene name PECAM1), CD34, von Willebrand Factor (vWF), tyrosine-protein kinase receptor Tie2 (TEK), VE-cadherin (CDH5), endoglin (ENG), CD146 (MCAM) and VEGFR2 (KDR)), astrocytes (blue, GFAP), immune cells (red; T-cells (CD3E), B-cells (CD19), all leucocytes (CD45, PTPRC), and monocytes/macrophages (F4/80, EMR1)), Müller glia cells (orange; aquaporin 4 (AQP4)), neurons (yellow; retinal ganglion cells (RNA binding fox-1 homolog 3, RBFOX3), amacrine cells (parvalbumin, PVALB), bipolar cells (PKC-α, PRKCA), horizontal cell (calbindin, CALB1), photoreceptors (rods, CD73 (NT5E); cones, transducing (GNAT1)), and retinal pigment epithelial cells (magenta; retinal pigment epithelium-specific 65 kDa protein (RPE65).

**Figure 2.**
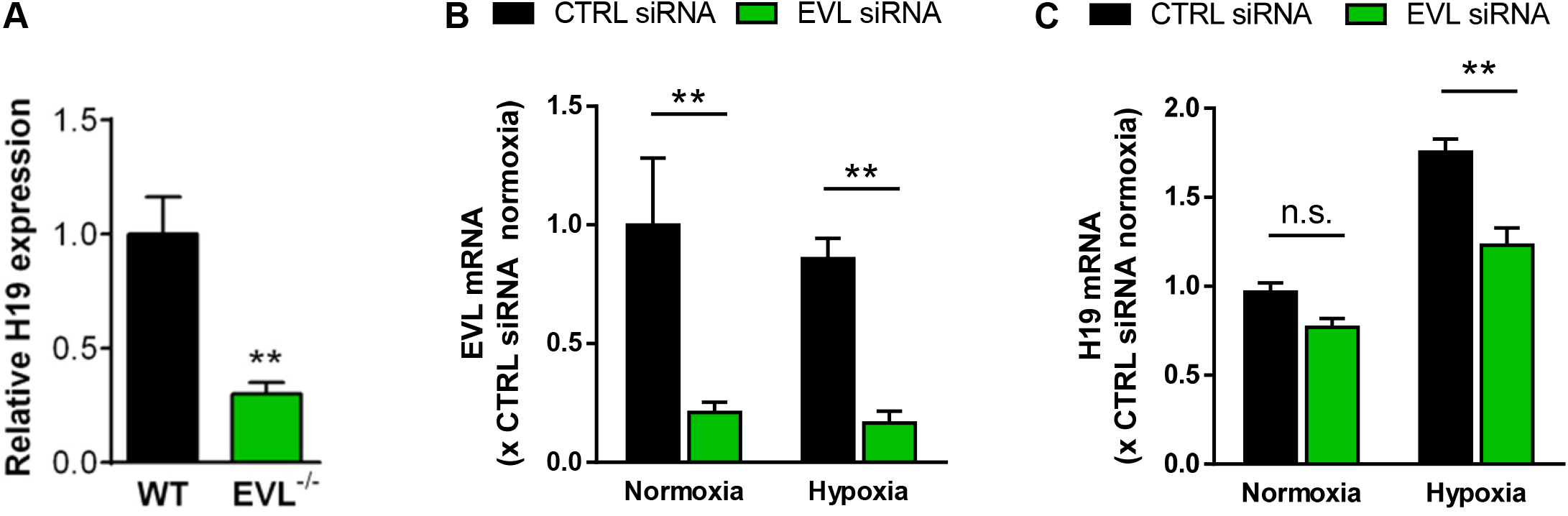
EVL-deficiency and siRNA-mediated knockdown impairs H19 expression in endothelial cells. **(A)** LncRNA H19 expression in CD31/CD34 double-positive endothelial cells from P5 EVL^-/-^ retinas, relative to wild-type controls, was assessed by RNA-sequencing as detailed in (A). **(B)** qRT-PCR analysis of siRNA-mediated knockdown of EVL in human umbilical vein endothelial cells after 14 hours under normal growth conditions (normoxia) or reduced oxygen levels (1% O_2_; hypoxia). **(C)** qRT-PCR analysis of H19 RNA levels in human umbilical vein endothelial cells subjected to siRNA-mediated knockdown of EVL after 14 hours under normoxic or hypoxic conditions. Error bars represent SEM, ^n.s.^P>0.05, **P<0.01, unpaired t-test.

**Figure 3.**
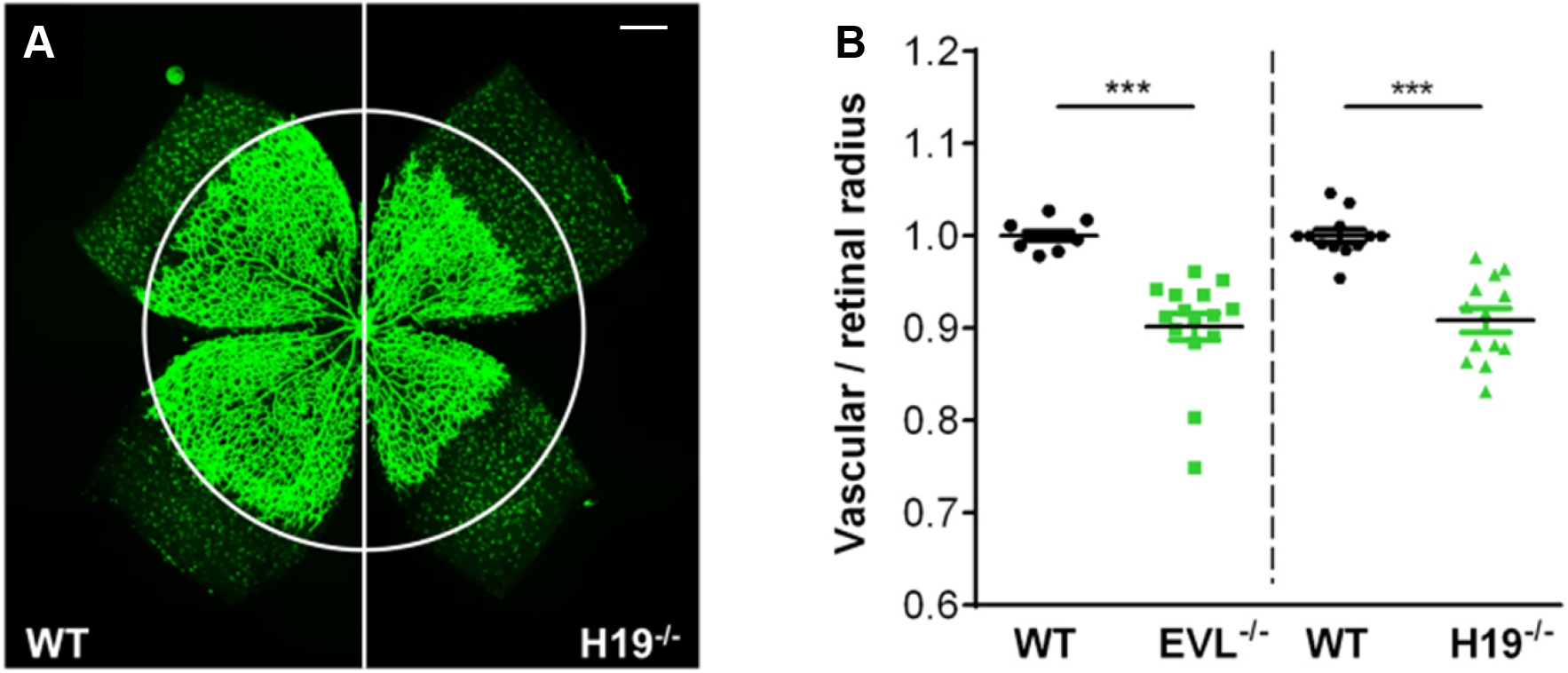
Delayed postnatal retinal angiogenesis in H19-deficient mice. **(A)** Eyes of wild-type and H19^-/-^ mice were harvested on P5, retinas isolated and the vasculature stained with Isolectin B4. **(B)** The radial outgrowth of the retinal vasculature was analyzed by confocal microscopy and normalized to the retinal radius as well as littermate controls. A comparison of the vascular spreading of H19- deficient animals to EVL^-/-^ mice revealed a delay in angiogenesis to a similar extent. Scale bar: 200 μm. H19^-/-^: n=12–14 animals, EVL^-/-^: n=10–12 animals of five litters per mouse line. Error bars represent SEM; *** p<0.001, one-way ANOVA with Bonferroni post test.

## Materials and methods

Retina whole mount staining and analysis. Global B6J-H19 deficient mice (H19^-/-^) (Hofmann et al., 2019) and wild-type littermate controls were sacrificed at P5 and eyeballs fixed in 4% PFA for 1 hour. Dissected retinas were equilibrated and permeabilized in PBlec buffer (1mM CaCl_2_, 1mM MgCl_2_, 0.1 mM MnCl_2_ and 0.5 % Triton X-100 in PBS, pH6.8) for 15 min and stained with FITC labelled Isolectin B4 (10 µg/ml, Sigma-Aldrich #L2895) in PBlec overnight at 4°C. After washing with PBS, retinas were flat-mounted in Mowiol mounting medium (25 % glycerol, 0.1 % Mowiol and 5 % DABCO in 0.1 M Tris, pH 8.5) or Dako fluorescence mounting medium and analyzed using a confocal microscope (Leica, SP8). The radial vascular outgrowth was assessed with a 10-fold objective by measuring the radius from the optic nerve to the vascular front and normalizing it to the retinal radius at 12 positions per retina and further normalizing it to the average of littermate controls. All experiments were approved by the governmental authorities. qRT-PCR analysis, human umbilical vein endothelial cells isolation, culture and transfection with siRNA was done as previously described (Zink et al., 2021).

## Acknowledgements

This work was supported by the Deutsche Forschungsgemeinschaft (SFB 834/A8 to PMB, SFB 834/A5 to IF, SFB834/B9 to RAB). PMB was also supported by the German Center for Cardiovascular Research (DZHK B14-028 SE). RAB was also supported by the European Research Council (‘NOVA’). The authors are indebted to Patrik Hofmann for scientific input and help with the H19^-/-^ mice.

